# Plant-PLMview: a database for identifying *cis*-regulatory sequences with preferential positions in gene-proximal regions of plants

**DOI:** 10.1101/2022.12.20.521192

**Authors:** Julien Rozière, Franck Samson, Cécile Guichard, Margot Correa, Sylvie Coursol, Marie-Laure Martin, Véronique Brunaud

## Abstract

**Background:** Establishing relationships between transcription factors and target genes is essential for understanding the mechanisms regulating gene expression, which play a fundamental role in plant adaptation to the local environment. Despite the importance of this research area and the tremendous progress in sequencing methods such as ChIP-seq and DAP-seq, we are still far from a complete reconstruction of the *cis*-regulatory landscape. Only a small number of transcription factors can be assessed from experimental data to identify their *cis*-regulatory binding sites. This highlights the role that *in silico* approaches can play to complement experimental data.

**Results:** We have developed Plant-PLMview, a web-accessible database for detecting preferentially *cis*-regulatory sequences in the gene-proximal regions of 20 plant species. Users of Plant-PLMview can (i) access the proximal regions of controlled genes from 20 plant species, (ii) query their own DNA motifs or access 840 *cis*-regulatory sequences from various plant resources, and (iii) use a tool called PLMdetect to search for preferentially located motifs in the gene-proximal regions of a list of genes. Results are displayed via a web interface with a list of DNA motifs preferentially located in a region near the start or end of genes, the distribution of these motifs and associated annotations. In addition, a graphical map of the preferential locations of the motifs in the 5’ and 3’-proximal regions of genes provides an overview of all motifs and genes examined.

**Conclusion:** Plant-PLMview provides the opportunity to study the proximal landscape of gene regulation in 20 plant genomes. The originality of the database lies in its ease of use thanks to the curated data (*cis*-regulatory sequences and proximal regions of genes) and the possibility to search through PLMdetect in the 5’-gene-proximal region, but also in the 3’-gene-proximal region, which is rarely explored. The web interface provides numerous graphical views that allow the users to get an overview and interpret the results more easily.

## BACKGROUND

The transcriptional activity of genes in plants is a fundamental process in the adaptation of these sessile organisms to the environment [1–6]. In this context, many combined experimental and *in silico* [7,8] efforts have been made to fully understand the mechanisms underlying the regulation of plant gene expression. Among the regulatory elements, transcription factors (TFs) play an important role in determining the tissue specificity and developmental stage specificity of gene expression [9].These proteins regulate the expression of one or more target genes by binding a very short DNA sequence, typically between 5 and 25 bases, in their enhancer or promoter regions.

The prediction and characterization of these DNA motifs or TF binding sites (TFBS) in a genome of several mega or giga bases presents an algorithmic and statistical challenge that can be partially solved by experimental approaches such as ChIP-seq or DAP-seq [10,11] to name the best known. These sequencing-based technologies have enabled extraordinary progress in the experimental determination of TFBSs, the results of which are stored in various databases such as JASPAR [12], REMAP [13] and Cis-BP [14]. These databases analyze the sequencing profile results of all these experiments to focus on TF-fixed regions and propose TFBS candidates, often using a position weight matrix (PWM) and logos. In this way, the last release of the database JASPAR [15] defined more than 600 TFBSs for the regulation of plant gene expression. However, these approaches have some limitations, such as the requirement of specific antibodies for each TF or TF family, the artificial accessibility of DNA in *in vitro* methods, and the large DNA regions to be studied, ranging from 100 to 10,000 bases [7]. To illustrate this limitation, only 372 of 2,000 TFs encoded by the plant model *Arabidopsis thaliana* were examined using ChIP-seq and DAP-seq to identify their binding sites (source: REMAP database). Fortunately, inference of TFBSs using *in silico* methods has become a relevant and complementary alternative. These methods can be rapidly and successfully applied to biological cross-constraints, such as experimental data and gene annotations. Interestingly, *cis*-regulatory sequences in proximal regions located in bases framing the transcription start site (TSS) or transcription termination site (TTS) appear to be associated with fixed topological constraints [16–18]. Consequently, these proximal regions are essentially rich in *cis*-regulatory sequences [19–21]. In this context, the *in silico* PLMdetect method [22,23] was developed to identify short DNA sequences that are overrepresented when the distance to the TSS or to the TTS is constrained, and are therefore referred to as preferentially located motifs (PLMs). Numerous applications are already showing interest in using PLMdetect to advance the characterization of the gene-proximal environment in targeted transcriptomic datasets and at the genome level for Arabidopsis and maize [21,24–28]. To make PLMdetect available to the entire community, we introduce Plant-PLMview, a web database that allows all users to explore the proximal environment of genes by performing PLM searches for 20 plant species. It has been simplified to limit the amount of input data, and interpretation of results is facilitated by visualization of PLMs. What makes Plant-PLMview unique is that it (i) provides the ability to work on a panel of 20 plant species selected for their representativeness of the angiosperm family, (ii) provides the ability to explore PLMs in the 3’-proximal region of genes, (iii) provides access to an extensive catalog of 840 *cis*-regulatory sequences, and (iv) provides a clear and interactive visualization of results.

## CONSTRUCTION AND CONTENT

### General structure

Plant-PLMview is an analysis database that is directly accessible via the Internet (http://plmview.ips2.universite-paris-saclay.fr). It is designed to identify *cis*-regulatory sequences in a range of genes of interest using PLMdetect, as described below. Plant-PLMview consists of two modules (Figure 1). The first module consists of two resources: (1) all 5’- and 3’-gene proximal sequences from 20 plant species and (2) the 840 *cis*-regulatory sequences integrated from experimental data. The second module is the query and tool part. It allows the user to run PLMdetect through the web interface by specifying a list of genes and a list of motifs. Once the analysis is completed, the results are returned by visualizing the PLMs on a map and as tables that can be downloaded.

**Figure 1:**
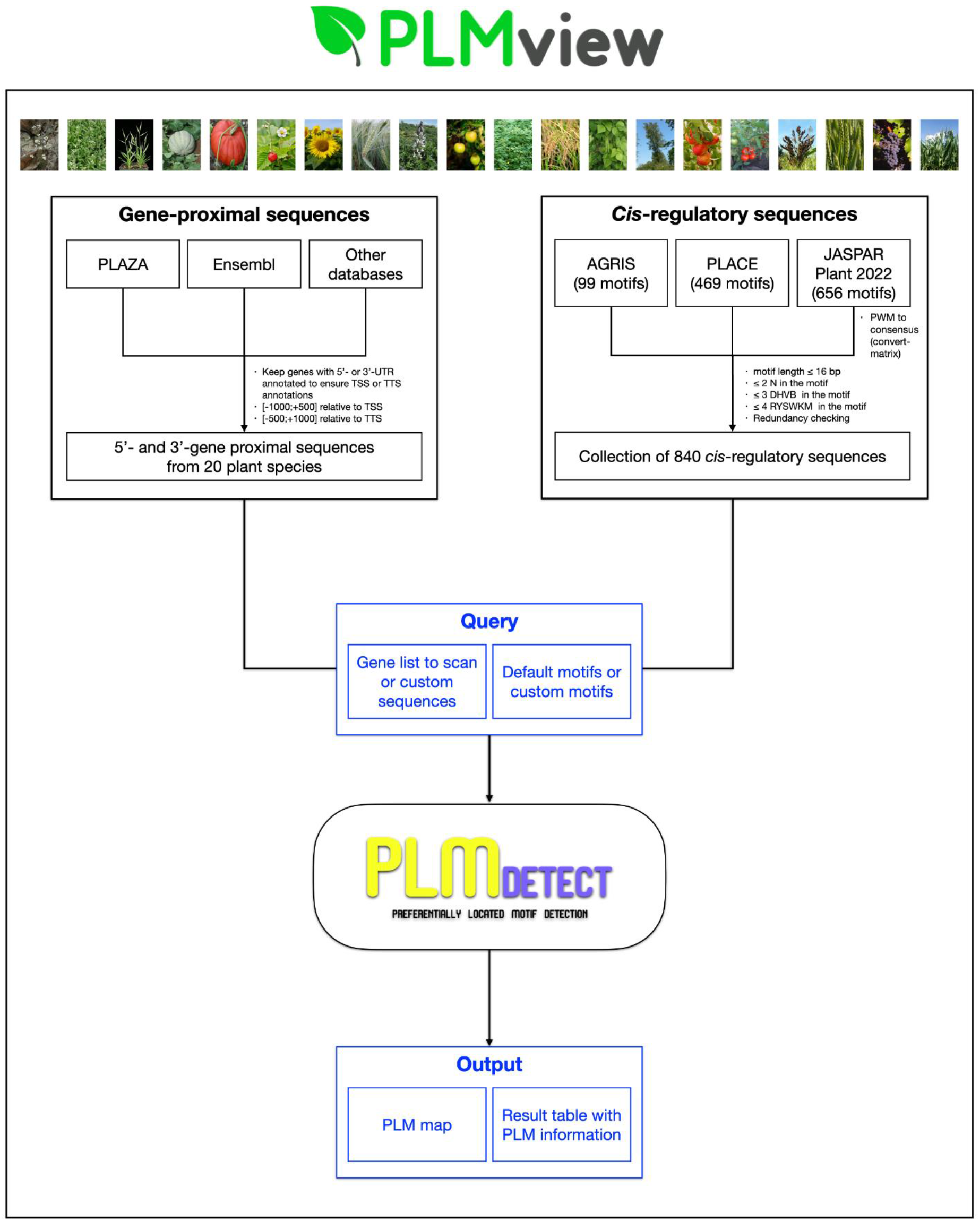
General structure of Plant-PLMview. The resource module is shown in black and the query module in blue.

### Description of PLMdetect implemented in Plant-PLMview

PLMdetect attempts to identify DNA motifs that are overrepresented at a given distance from the TSS or the TTS. Given a set of sequences and a motif, PLMdetect counts the number of occurrences of the motif at each position in the sequence set to obtain a motif distribution. Then, a linear regression is estimated for the neutral region corresponding to the first 500 bases of the sequences, and the predicted values with an associated 99% confidence interval are calculated for the region under study corresponding to the remaining sequence downstream of the neutral region. Finally, PLMdetect declares the motif as PLM if a peak in the region under investigation exceeds the confidence interval.

Each PLM is characterized by three indicators: (i) a preferential position, defined as the position of the maximum distribution peak, (ii) a functional window corresponding to the boundaries of the peak, and (iii) a score determined by the peak height (Figure 2).

**Figure 2:**
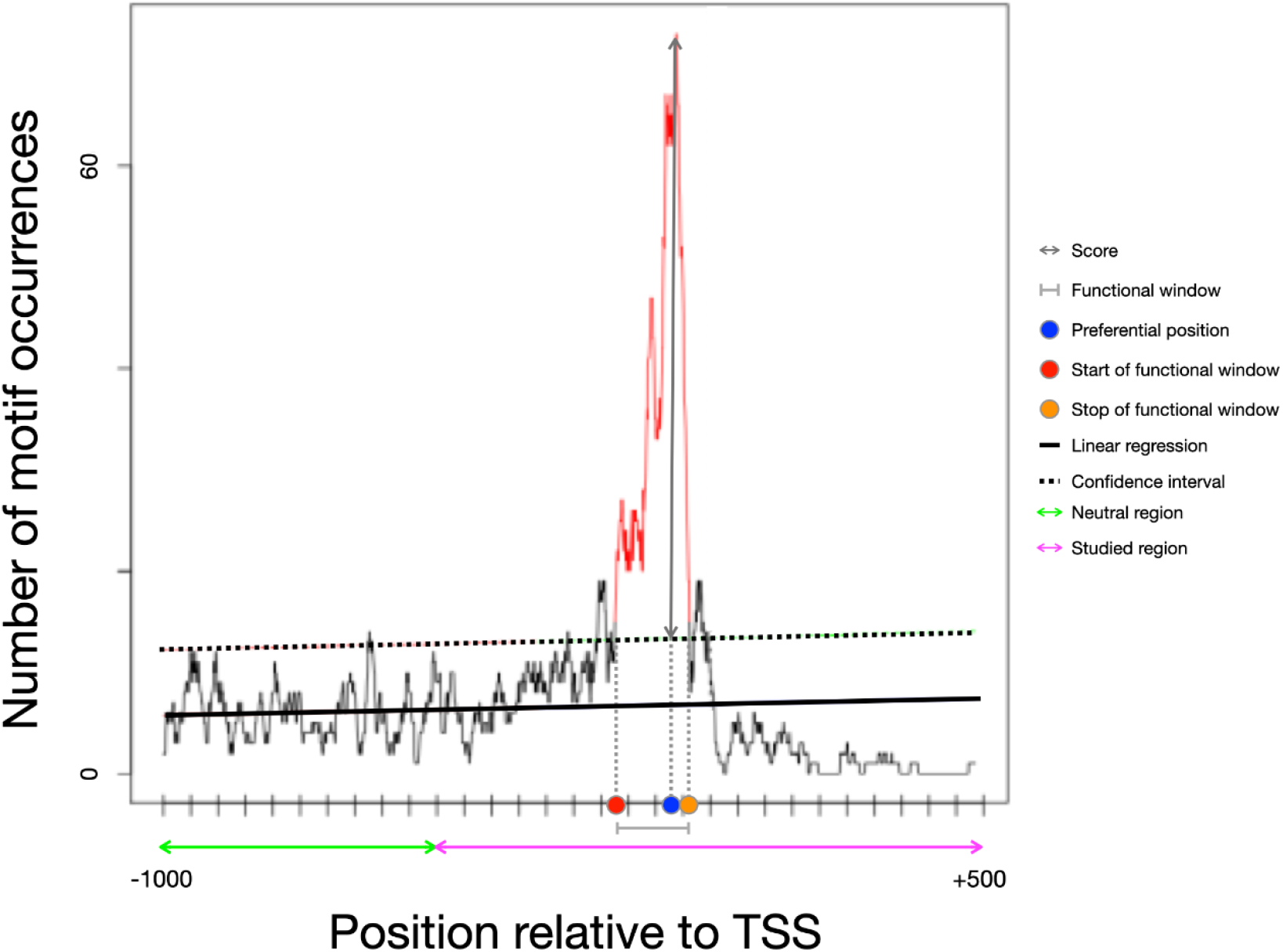
Example of a motif distribution illustrating a PLM and its characteristics. This graph represents the MCACGTGGC distribution in a subset of *A. thaliana* 5’-gene-proximal regions. In this example, the distribution is computed on the [-1000;+500] bp interval relative to the TSS.

### Gene-proximal sequence resource

Plant-PLMview contains gene-proximal sequences of 20 plant species covering the diversity of the angiosperm family with representatives of the Brassicaceae, Rosaceae, Fabaceae, Cucurbitaceae, Solanaceae, and the Poaceae. These plant genomes and annotations were mainly obtained from the Phytozome [29] and Ensembl Plant [30] databases (Table 1), and only gene sequences with an annotated 5’ (3’)-UTR were considered to ensure the TSS (TTS) annotations and thus the quality of the detected PLM. The sequences included in the database consist of the interval [-1000;+500] relative to TSS for the 5’-gene-proximal region and the interval [-500;+1000] relative to TTS for the 3’-gene-proximal region. The sequences of the antisense genes were added in reverse order to standardize the application of PLMdetect.

**Table 1:**
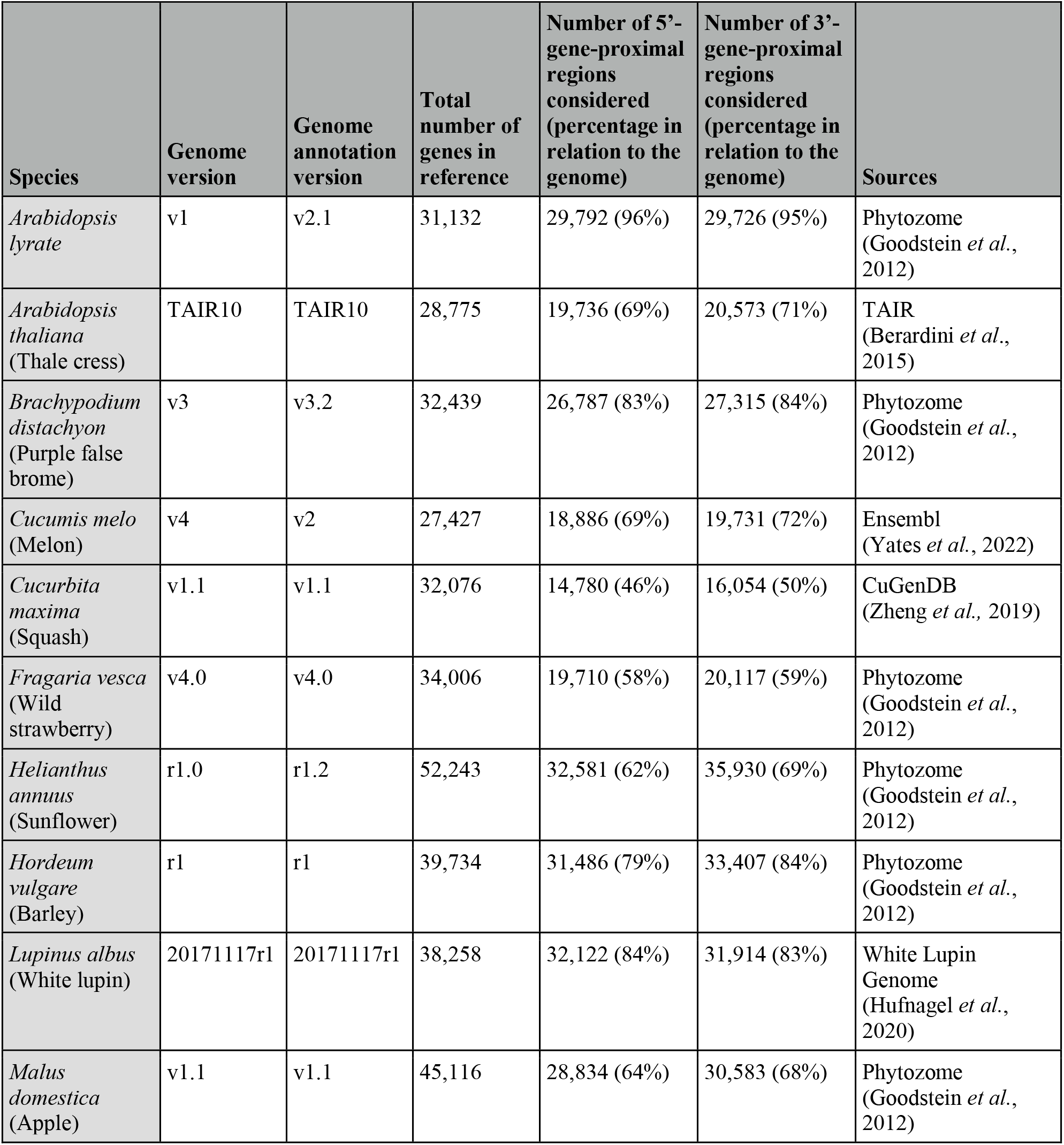
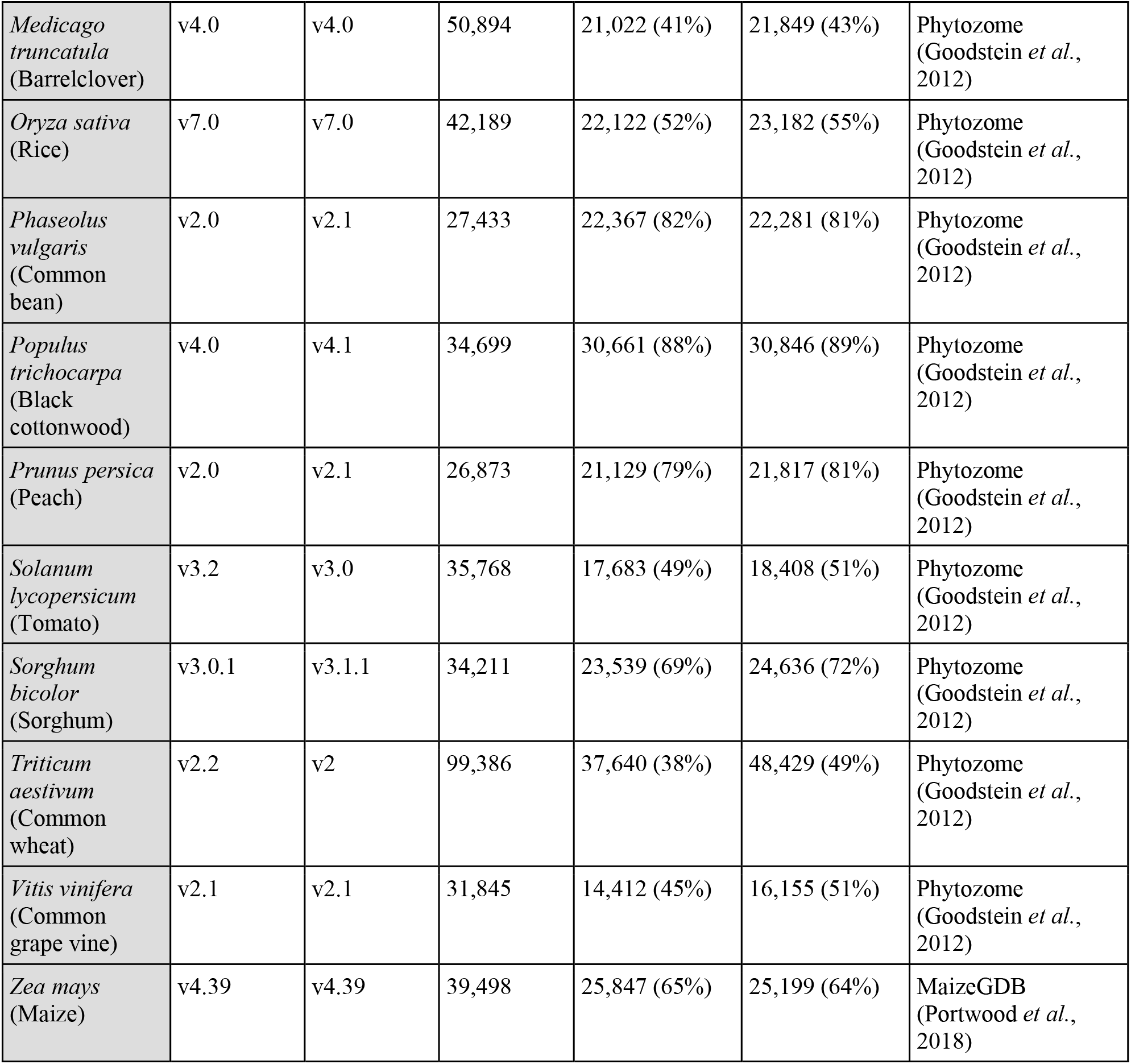
Description of the 20 plant species available in Plant-PLMview. Indicated are the genome versions, annotation versions, sources, number of genes described in the species, and genes included in the analyses of the 5’ and 3’-gene proximal regions.

| Species | Genome version | Genome annotation version | Total number of genes in reference | Number of 5'-gene-proximal regions considered (percentage in relation to the genome) | Number of 3'-gene-proximal regions considered (percentage in relation to the genome) | Sources |
| --- | --- | --- | --- | --- | --- | --- |
| *Arabidopsis lyrata* | v1 | v2.1 | 31,132 | 29,792 (96%) | 29,726 (95%) | Phytozome (Goodstein *et al.*, 2012) |
| *Arabidopsis thaliana* (Thale cress) | TAIR10 | TAIR10 | 28,775 | 19,736 (69%) | 20,573 (71%) | TAIR (Berardini *et al.*, 2015) |
| *Brachypodium distachyon* (Purple false brome) | v3 | v3.2 | 32,439 | 26,787 (83%) | 27,315 (84%) | Phytozome (Goodstein *et al.*, 2012) |
| *Cucumis melo* (Melon) | v4 | v2 | 27,427 | 18,886 (69%) | 19,731 (72%) | Ensembl (Yates *et al.*, 2022) |
| *Cucurbita maxima* (Squash) | v1.1 | v1.1 | 32,076 | 14,780 (46%) | 16,054 (50%) | CuGenDB (Zheng *et al.*, 2019) |
| *Fragaria vesca* (Wild strawberry) | v4.0 | v4.0 | 34,006 | 19,710 (58%) | 20,117 (59%) | Phytozome (Goodstein *et al.*, 2012) |
| *Helianthus annuus* (Sunflower) | r1.0 | r1.2 | 52,243 | 32,581 (62%) | 35,930 (69%) | Phytozome (Goodstein *et al.*, 2012) |
| *Hordeum vulgare* (Barley) | r1 | r1 | 39,734 | 31,486 (79%) | 33,407 (84%) | Phytozome (Goodstein *et al.*, 2012) |
| *Lupinus albus* (White lupin) | 20171117r1 | 20171117r1 | 38,258 | 32,122 (84%) | 31,914 (83%) | White Lupin Genome (Hufnagel *et al.*, 2020) |
| *Malus domestica* (Apple) | v1.1 | v1.1 | 45,116 | 28,834 (64%) | 30,583 (68%) | Phytozome (Goodstein *et al.*, 2012) |
| *Medicago truncatula*<br>(Barrelclover) | v4.0 | v4.0 | 50,894 | 21,022 (41%) | 21,849 (43%) | Phytozome<br>(Goodstein *et al.*,<br>2012) |
| *Oryza sativa*<br>(Rice) | v7.0 | v7.0 | 42,189 | 22,122 (52%) | 23,182 (55%) | Phytozome<br>(Goodstein *et al.*,<br>2012) |
| *Phaseolus vulgaris*<br>(Common bean) | v2.0 | v2.1 | 27,433 | 22,367 (82%) | 22,281 (81%) | Phytozome<br>(Goodstein *et al.*,<br>2012) |
| *Populus trichocarpa*<br>(Black cottonwood) | v4.0 | v4.1 | 34,699 | 30,661 (88%) | 30,846 (89%) | Phytozome<br>(Goodstein *et al.*,<br>2012) |
| *Prunus persica*<br>(Peach) | v2.0 | v2.1 | 26,873 | 21,129 (79%) | 21,817 (81%) | Phytozome<br>(Goodstein *et al.*,<br>2012) |
| *Solanum lycopersicum*<br>(Tomato) | v3.2 | v3.0 | 35,768 | 17,683 (49%) | 18,408 (51%) | Phytozome<br>(Goodstein *et al.*,<br>2012) |
| *Sorghum bicolor*<br>(Sorghum) | v3.0.1 | v3.1.1 | 34,211 | 23,539 (69%) | 24,636 (72%) | Phytozome<br>(Goodstein *et al.*,<br>2012) |
| *Triticum aestivum*<br>(Common wheat) | v2.2 | v2 | 99,386 | 37,640 (38%) | 48,429 (49%) | Phytozome<br>(Goodstein *et al.*,<br>2012) |
| *Vitis vinifera*<br>(Common grape vine) | v2.1 | v2.1 | 31,845 | 14,412 (45%) | 16,155 (51%) | Phytozome<br>(Goodstein *et al.*,<br>2012) |
| *Zea mays*<br>(Maize) | v4.39 | v4.39 | 39,498 | 25,847 (65%) | 25,199 (64%) | MaizeGDB<br>(Portwood *et al.*,<br>2018) |

### *Cis*-regulatory sequence resource

Plant-PLMview integrates experimentally characterized *cis*-regulatory sequences from four complementary plant-related resources: (i) 469 sequences from PLACE[31], containing all sequences described up to 2007, (ii) 99 sequences from AGRIS[32] for *A. thaliana*, (iii) 15 sequences from the work of Bernard *et al*. [23] corresponding to TATA-like sequences, and (iv) 656 sequences from JASPAR [33] plant part 2022 listing TFBSs identified mainly from PBM, ChIP-seq, DAP-seq or SELEX-seq data. Since JASPAR stores the information as PWMs, they have to be converted into consensus sequences. This was done using the convert-matrix tool of the RSAT suite [34] with default parameters.

We eliminated redundancies between resources and removed sequences larger than 16 bp or with too many IUPAC indeterminacies (≥ 2 N, ≥ 3 DHVB, ≥ 4 RYSWKM). After this curation, the 840 unique *cis*-regulatory sequences were organized into a motif-oriented resource. It is possible to explore this sequence catalog from the query page using the “access to motif catalog” button.

We chose a hierarchical graph to highlight the overlaps and inclusion links between the different *cis*-regulatory sequences. Figure 3 shows the screenshot of the motif resource when a query was submitted to TGACG. The information provided is the description of the sequence, its other names, any associated bibliographic references, the list of species in which the sequence has been described, and the three main source databases (PLACE, AGRIS, JASPAR).

**Figure 3:**
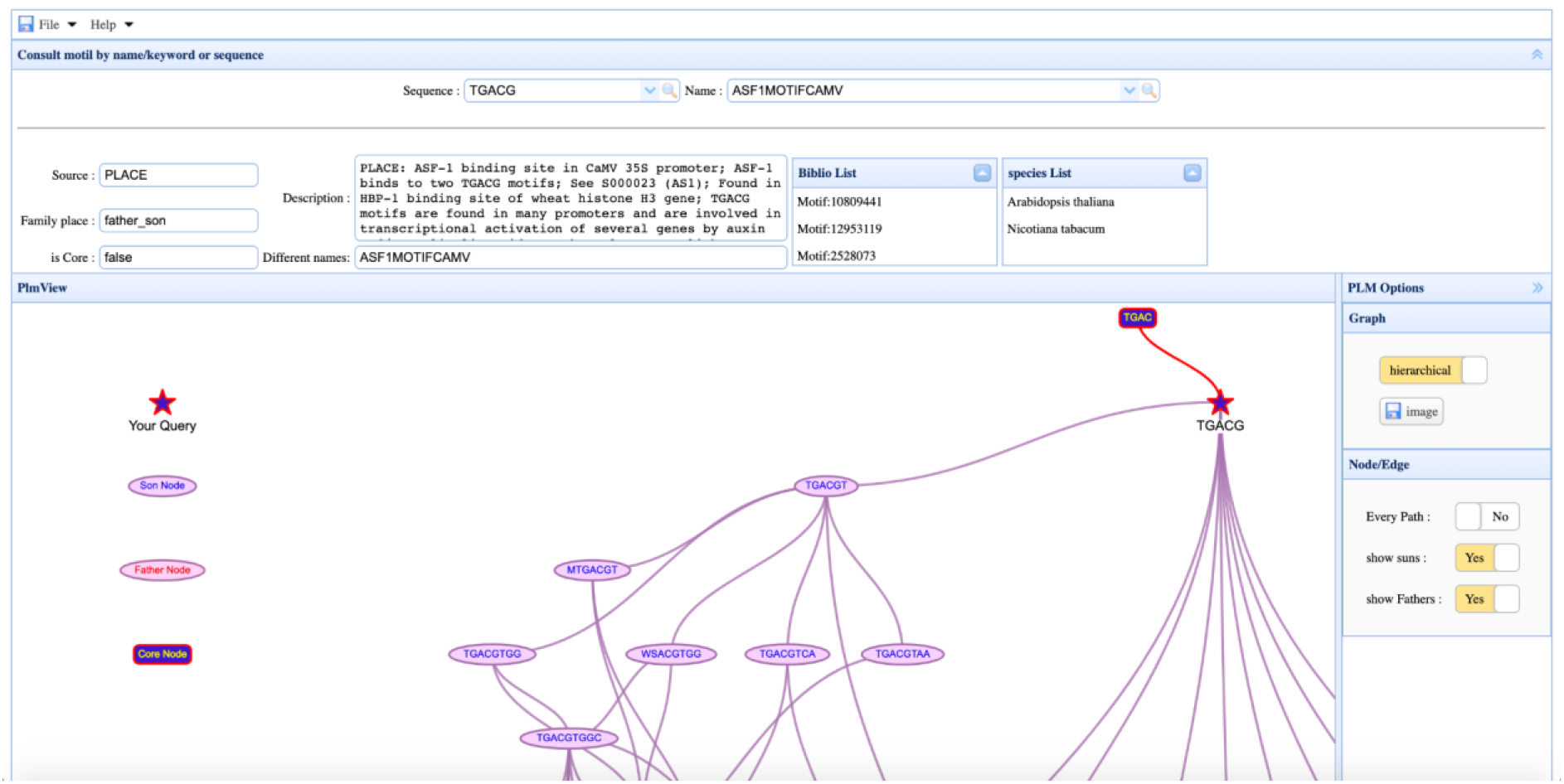
TGACG family visualization and information. The motif-resource interface shows the details of each *cis*-regulatory sequence in our catalog. It provides access to the description of the motif, its source database, its place in the family, other names, associated bibliographic references, and the species in which this sequence was described. A sequence family is defined by a set of sequences grouped by inclusion links. In a given family, if sequence 1 is included in sequence 2, sequence 1 is considered as the father of sequence 2. Conversely, sequence 2 is the son of sequence 1. In this example, TGACG is the father of TGACGT and the son of TGAC. The graphical part allows the user to visualize the family of the sequence and access each member by clicking on it.

### Database and web implementation

The database is built on the relational system PostgreSQL (version 13). Plant-PLMview allows downloading the list of 840 motifs and the studied 5’ and 3’ proximal regions as Fasta files. The interfaces of Plant-PLMview are written in Javascript Python3 and use the modules Flask, psycopg2, easyui, vis, and d3js.

## UTILITY AND DISCUSSION

The interface is typically used to identify *cis*-regulatory sequences among differentially expressed genes or co-expressed genes.

### Use Case

We illustrate this by analyzing a group of genes from Bueso *et al*. [25], the example accessible via the “DEMO” button on the query page. First, the user must select a species, enter the gene identifiers and the gene-proximal region (5’ or 3’) to be analyzed. In this use case, the selected species is *A. thaliana*, the queried genes are specified in the “paste a list of gene IDs” and we want to examine the 5’-gene proximal region (Figure 4). After entering his email address, the user clicks on the “Run Demo’’ button to get a link with all the results. For further analysis, the user clicks the “Run PLM” button after entering the inputs.

**Figure 4:**
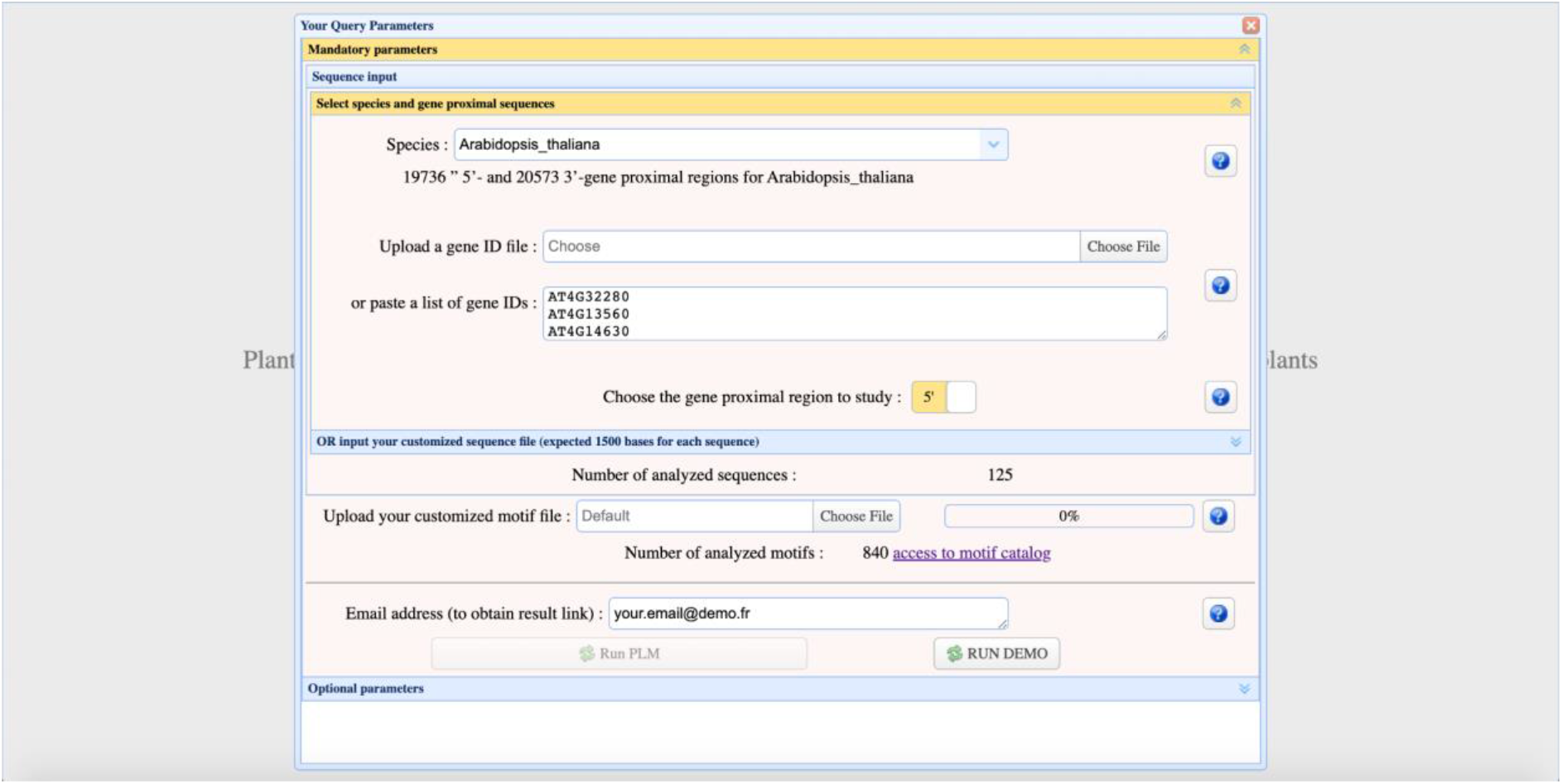
Plant-PLMview query interface. To start a query, the user must select a species, specify the gene IDs, and select the gene-proximal region to be examined.

For advanced users, the interface also provides optional parameters for filtering PLMs by score, setting the size of the sliding window for counting *cis*-regulatory sequences, and completing the search in a directed manner by disabling the “Two strands” option. In the use case, we leave the optional parameters as default. After some time (depending on the sequence and motif number entered), Plant-PLMview returns a map of all identified PLMs. It allows the user to see at a glance the *cis*-regulatory motif candidates and their position in the gene-proximal region. This map is interactive: it is possible to zoom in and out, filter by genes or PLMs, and take screenshots. In our use case, Plant-PLMview identified 40 PLMs distributed across 125 genes (Figure 5). The map shows a strong presence of TATA-box motifs and their TATA-box derivatives upstream of TSS of the gene tested. It also shows strong conservation of some PLMs downstream of TSS, such as TGACG and GANTTNC, which are known to be bound by bZIP and MYB TFs, respectively. The visualization proposed by Plant-PLMview also highlights combinations of PLMs that might be involved in this co-regulation of genes. Taking TGACG and GANTTNC PLMs as an example, it shows that these PLMs are co-present in 31 genes, suggesting a potential combined effect of bZIP and MYB TFs in these 5’-gene proximal regions.

**Figure 5:**
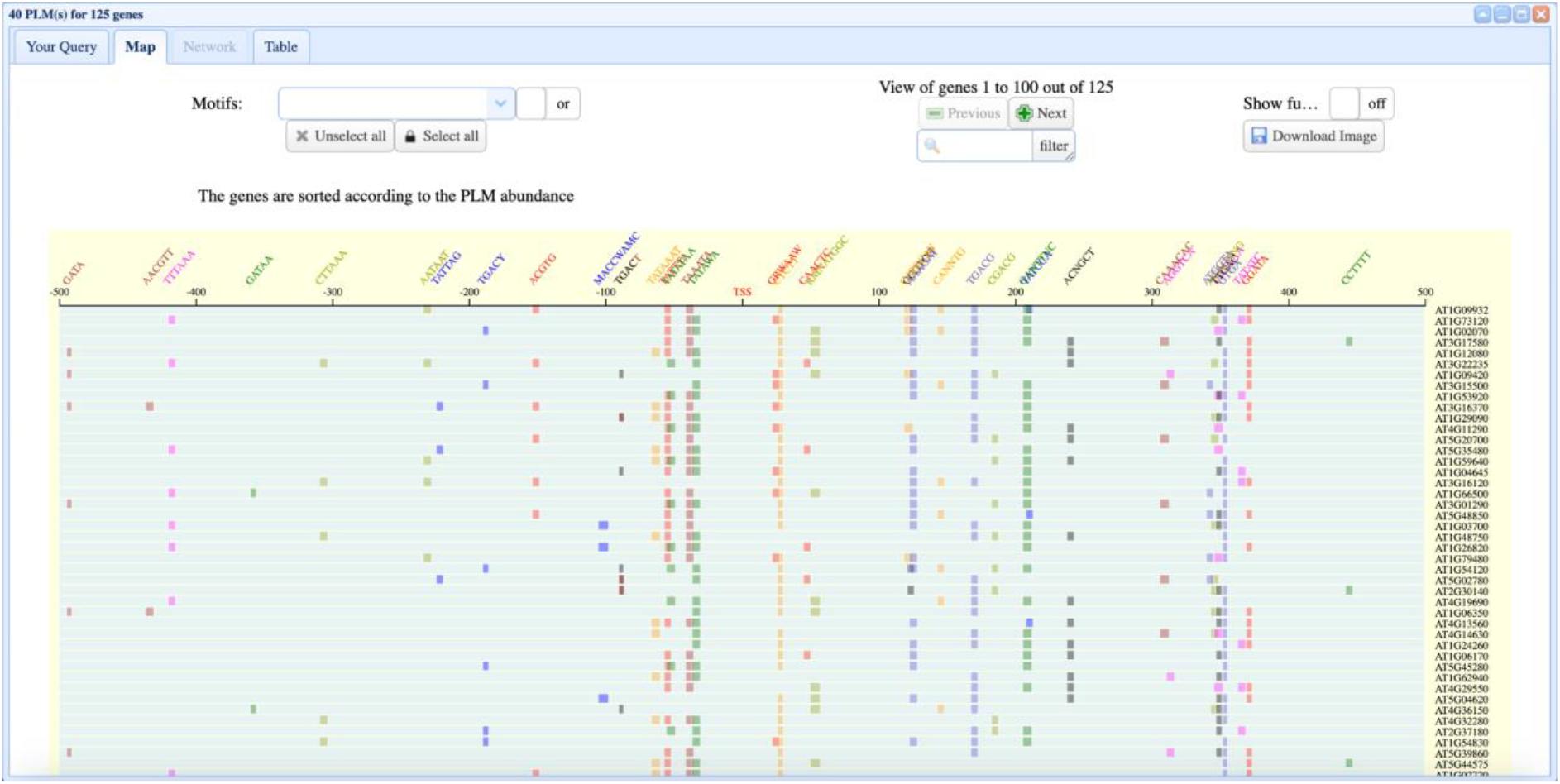
Example of a PLM map. The map shows the PLMs identified by gene. The map is interactive and can be modified by filtering genes or PLMs. The visualization can be downloaded by clicking on the “Download Image” button.

The results are also reflected in a table that contains all the information for each PLM (Figure 6). This includes the DNA motif, the distribution of its occurrence, its score, its positional window, its preferential position, its function, and the known information about this *cis*-regulatory sequence. This table is downloaded using the “Download Results’’ button. The results are then organized in different directories: the “PLM_lists” directory contains for each PLM the list of genes in which it occurs, the “PLM_graphs’’ directory contains the distribution of the occurrence and a file with all available information about these PLMs.

**Figure 6:**
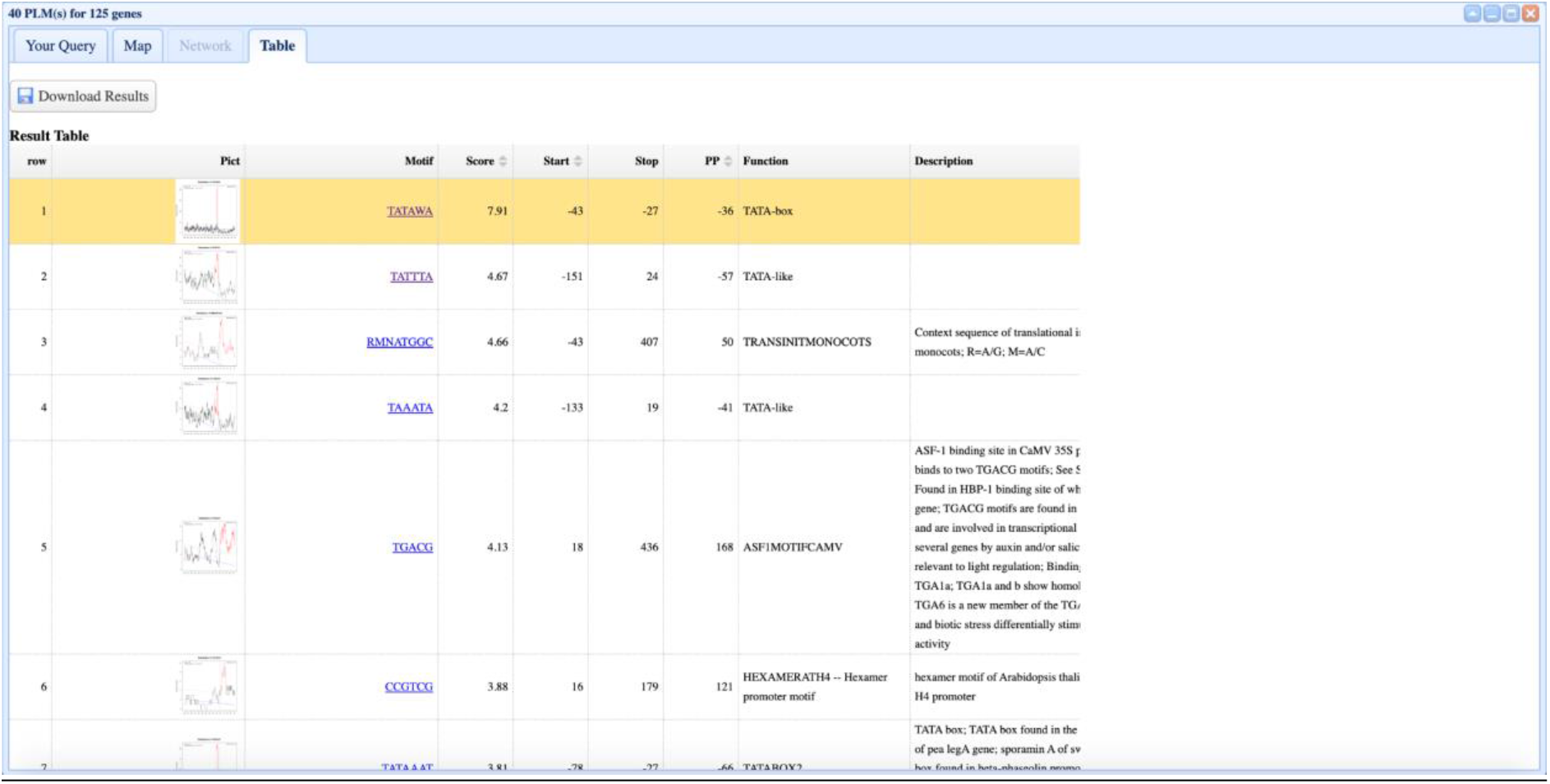
Example of a PLM result table. This table lists the distribution of motifs for each PLM, the positional data, and the functions known in the bibliography. It is possible to click on the motif of each PLM to see it in the motif resource.

To locate the PLM with respect to the *cis*-regulatory motifs stored in Plant-PLMview, the user simply clicks on the name of the motif in the results table and an image appears as in Figure 3.

### Future implementations

Plant-PLMview allows users to identify candidate *cis*-regulatory sequences underlying potential co-regulation of gene groups belonging to any of the 20 available species among the 840 *cis*-regulatory sequences in the database. For users who wish to examine species or *cis*-regulatory sequences not included in Plant-PLMview, we have provided the ability to provide 5’- or 3’-gene-proximal regions and custom *cis*-regulatory sequences. The user-provided gene-proximal region file should list sequences defined by the range [-1000;+500] bp relative to TSS and [-500;+1000] bp relative to TTS. The sequence orientation guidelines given in the “Genomes and gene-proximal sequences’’ section should also be followed. The custom motif file must contain DNA motifs that follow the IUPAC codes and are separated by a line break. The provided motifs must comply with the filter guidelines (if motif size and indeterminacies are not followed, the motif will be removed) in the “*Cis*-regulatory sequence data sources and processing” section. In the current version of Plant-PLMview, only one species can be queried at a time. Our near-term goal is to provide access to a cross-species PLM search to investigate coevolution of *cis*-regulatory motifs by providing the ability to select multiple species simultaneously on the query page and enter gene identifiers.

## CONCLUSIONS

The search for *cis*-regulatory sequences involved in the regulation of gene clusters of interest is a widely studied topic. Access to *in silico* approaches remains limited for laypersons, indicating the need for user-friendly tools to accelerate research in this area. Here, we introduced Plant-PLMview, which provides an easy-to-use web interface and easy-to-interpret results. This is made possible by the numerous pre-processing steps of the data and PLM visualization as an interactive map.

## DECLARATIONS

### Ethics approval and consent to participate

‘Not applicable’

### Consent for publication

‘Not applicable’

### Availability of data and materials

Database access URL: http://plmview.ips2.universite-paris-saclay.fr

All the proximal gene regions from 20 plant species of PLMview database, are extracted from resources described in table 1 and accessible for download by web link http://tools.ips2.u-psud.fr/ftp/PLMview-data

### Authors’ contributions

All the authors conceived the project. JR and VB supervised the project and extracted resources data. CG created the database and integrated data. FS developed the web interface. MC and M-LM tested the database. JR, VB, M-LM and SC wrote the manuscript. All the authors read the paper and approved the submitted version.

## Acknowledgements

We thank F. Desprez for installing and managing the system and server at the IPS2 Institute.

## REFERENCES

1. Yocca, A.E.; Edger, P.P. Current Status and Future Perspectives on the Evolution of Cis-Regulatory Elements in Plants. Current Opinion in Plant Biology 2022, 65, 102139, doi:10.1016/j.pbi.2021.102139.

2. Alonge, M.; Wang, X.; Benoit, M.; Soyk, S.; Pereira, L.; Zhang, L.; Suresh, H.; Ramakrishnan, S.; Maumus, F.; Ciren, D.; et al. Major Impacts of Widespread Structural Variation on Gene Expression and Crop Improvement in Tomato. Cell 2020, 182, 145-161.e23, doi:10.1016/j.cell.2020.05.021.

3. Azodi, C.B.; Lloyd, J.P.; Shiu, S.-H. The Cis-Regulatory Codes of Response to Combined Heat and Drought Stress in Arabidopsis Thaliana. NAR Genomics and Bioinformatics 2020, 2, doi:10.1093/nargab/lqaa049.

4. Waters, A.J.; Makarevitch, I.; Noshay, J.; Burghardt, L.T.; Hirsch, C.N.; Hirsch, C.D.; Springer, N.M. Natural Variation for Gene Expression Responses to Abiotic Stress in Maize. The Plant Journal 2017, 89, 706–717, doi:10.1111/tpj.13414.

5. Zhou, P.; Enders, T.A.; Myers, Z.A.; Magnusson, E.; Crisp, P.A.; Noshay, J.M.; Gomez-Cano, F.; Liang, Z.; Grotewold, E.; Greenham, K.; et al. Prediction of Conserved and Variable Heat and Cold Stress Response in Maize Using Cis-Regulatory Information. The Plant Cell 2022, 34, 514–534, doi:10.1093/plcell/koab267.

6. Liu, S.; Li, C.; Wang, H.; Wang, S.; Yang, S.; Liu, X.; Yan, J.; Li, B.; Beatty, M.; Zastrow-Hayes, G.; et al. Mapping Regulatory Variants Controlling Gene Expression in Drought Response and Tolerance in Maize. Genome Biology 2020, 21, 163, doi:10.1186/s13059-020-02069-1.

7. Lai, X.; Stigliani, A.; Vachon, G.; Carles, C.; Smaczniak, C.; Zubieta, C.; Kaufmann, K.; Parcy, F. Building Transcription Factor Binding Site Models to Understand Gene Regulation in Plants. Mol Plant 2018, doi:10.1016/j.molp.2018.10.010.

8. Boeva, V. Analysis of Genomic Sequence Motifs for Deciphering Transcription Factor Binding and Transcriptional Regulation in Eukaryotic Cells. Frontiers in Genetics 2016, 7.

9. Song, L.; Huang, S.C.; Wise, A.; Castanon, R.; Nery, J.R.; Chen, H.; Watanabe, M.; Thomas, J.; Bar-Joseph, Z.; Ecker, J.R. A Transcription Factor Hierarchy Defines an Environmental Stress Response Network. Science 2016, 354, aag1550, doi:10.1126/science.aag1550.

10. Jayaram, N.; Usvyat, D.; R. Martin, A.C. Evaluating Tools for Transcription Factor Binding Site Prediction. BMC Bioinformatics 2016, doi:10.1186/s12859-016-1298-9.

11. Bartlett, A.; O’Malley, R.C.; Huang, S.C.; Galli, M.; Nery, J.R.; Gallavotti, A.; Ecker, J.R. Mapping Genome-Wide Transcription Factor Binding Sites Using DAP-Seq. Nat Protoc 2017, 12, 1659–1672, doi:10.1038/nprot.2017.055.

12. Fornes, O.; Castro-Mondragon, J.A.; Khan, A.; van der Lee, R.; Zhang, X.; Richmond, P.A.; Modi, B.P.; Correard, S.; Gheorghe, M.; Baranašić, D.; et al. JASPAR 2020: Update of the Open-Access Database of Transcription Factor Binding Profiles. Nucleic Acids Res 2020, 48, D87–D92, doi:10.1093/nar/gkz1001.

13. Chèneby, J.; Ménétrier, Z.; Mestdagh, M.; Rosnet, T.; Douida, A.; Rhalloussi, W.; Bergon, A.; Lopez, F.; Ballester, B. ReMap 2020: A Database of Regulatory Regions from an Integrative Analysis of Human and Arabidopsis DNA-Binding Sequencing Experiments. Nucleic Acids Research 2020, 48, D180–D188, doi:10.1093/nar/gkz945.

14. Weirauch, M.T.; Yang, A.; Albu, M.; Cote, A.G.; Montenegro-Montero, A.; Drewe, P.; Najafabadi, H.S.; Lambert, S.A.; Mann, I.; Cook, K.; et al. Determination and Inference of Eukaryotic Transcription Factor Sequence Specificity. Cell 2014, 158, 1431–1443, doi:10.1016/j.cell.2014.08.009.

15. Castro-Mondragon, J.A.; Riudavets-Puig, R.; Rauluseviciute, I.; Lemma, R.B.; Turchi, L.; Blanc-Mathieu, R.; Lucas, J.; Boddie, P.; Khan, A.; Manosalva Pérez, N.; et al. JASPAR 2022: The 9th Release of the Open-Access Database of Transcription Factor Binding Profiles. Nucleic Acids Res 2022, 50, D165–D173, doi:10.1093/nar/gkab1113.

16. Yamamoto, Y.Y.; Ichida, H.; Matsui, M.; Obokata, J.; Sakurai, T.; Satou, M.; Seki, M.; Shinozaki, K.; Abe, T. Identification of Plant Promoter Constituents by Analysis of Local Distribution of Short Sequences. BMC Genomics 2007, 8, 67, doi:10.1186/1471-2164-8-67.

17. Bernardes, W.S.; Menossi, M. Plant 3’ Regulatory Regions From MRNA-Encoding Genes and Their Uses to Modulate Expression. Front Plant Sci 2020, 11, 1252, doi:10.3389/fpls.2020.01252.

18. Jores, T.; Tonnies, J.; Dorrity, M.W.; Cuperus, J.T.; Fields, S.; Queitsch, C. Identification of Plant Enhancers and Their Constituent Elements by STARR-Seq in Tobacco Leaves[OPEN]. Plant Cell 2020, 32, 2120–2131, doi:10.1105/tpc.20.00155.

19. Yu, C.-P.; Lin, J.-J.; Li, W.-H. Positional Distribution of Transcription Factor Binding Sites in Arabidopsis Thaliana. Scientific Reports 2016, 6, 25164, doi:10.1038/srep25164.

20. Ksouri, N.; Castro-Mondragón, J.A.; Montardit-Tarda, F.; van Helden, J.; Contreras-Moreira, B.; Gogorcena, Y. Tuning Promoter Boundaries Improves Regulatory Motif Discovery in Nonmodel Plants: The Peach Example. Plant Physiol 2021, 185, 1242–1258, doi:10.1093/plphys/kiaa091.

21. Rozière, J.; Guichard, C.; Brunaud, V.; Martin, M.-L.; Coursol, S. A Comprehensive Map of Preferentially Located Motifs Reveals Distinct Proximal Cis-Regulatory Sequences in Plants. Front Plant Sci 2022, 13, 976371, doi:10.3389/fpls.2022.976371.

22. Bernard, V.; Lecharny, A.; Brunaud, V. Improved Detection of Motifs with Preferential Location in Promoters. Genome 2010, 53, 739–752, doi:10.1139/g10-042.

23. Bernard, V.; Brunaud, V.; Lecharny, A. TC-Motifs at the TATA-Box Expected Position in Plant Genes: A Novel Class of Motifs Involved in the Transcription Regulation. BMC Genomics 2010, 11, 166, doi:10.1186/1471-2164-11-166.

24. Frei dit Frey, N.; Garcia, A.V.; Bigeard, J.; Zaag, R.; Bueso, E.; Garmier, M.; Pateyron, S.; de Tauzia-Moreau, M.-L.; Brunaud, V.; Balzergue, S.; et al. Functional Analysis of Arabidopsis Immune-Related MAPKs Uncovers a Role for MPK3 as Negative Regulator of Inducible Defences. Genome Biol. 2014, 15, R87, doi:10.1186/gb-2014-15-6-r87.

25. Bueso, E.; Muñoz-Bertomeu, J.; Campos, F.; Brunaud, V.; Martínez, L.; Sayas, E.; Ballester, P.; Yenush, L.; Serrano, R. ARABIDOPSIS THALIANA HOMEOBOX25 Uncovers a Role for Gibberellins in Seed Longevity. Plant Physiol. 2014, 164, 999–1010, doi:10.1104/pp.113.232223.

26. Bueso, E.; Muñoz-Bertomeu, J.; Campos, F.; Martínez, C.; Tello, C.; Martínez-Almonacid, I.; Ballester, P.; Simón-Moya, M.; Brunaud, V.; Yenush, L.; et al. Arabidopsis COGWHEEL1 Links Light Perception and Gibberellins with Seed Tolerance to Deterioration. Plant J. 2016, 87, 583–596, doi:10.1111/tpj.13220.

27. Cuello, C.; Baldy, A.; Brunaud, V.; Joets, J.; Delannoy, E.; Jacquemot, M.-P.; Botran, L.; Griveau, Y.; Guichard, C.; Soubigou-Taconnat, L.; et al. A Systems Biology Approach Uncovers a Gene Co-Expression Network Associated with Cell Wall Degradability in Maize. PLoS One 2019, 14, e0227011, doi:10.1371/journal.pone.0227011.

28. Del Prete, S.; Molitor, A.; Charif, D.; Bessoltane, N.; Soubigou-Taconnat, L.; Guichard, C.; Brunaud, V.; Granier, F.; Fransz, P.; Gaudin, V. Extensive Nuclear Reprogramming and Endoreduplication in Mature Leaf during Floral Induction. BMC Plant Biology 2019, 19, 135, doi:10.1186/s12870-019-1738-6.

29. Goodstein, D.M.; Shu, S.; Howson, R.; Neupane, R.; Hayes, R.D.; Fazo, J.; Mitros, T.; Dirks, W.; Hellsten, U.; Putnam, N.; et al. Phytozome: A Comparative Platform for Green Plant Genomics. Nucleic Acids Research 2012, 40, D1178–D1186, doi:10.1093/nar/gkr944.

30. Yates, A.D.; Allen, J.; Amode, R.M.; Azov, A.G.; Barba, M.; Becerra, A.; Bhai, J.; Campbell, L.I.; Carbajo Martinez, M.; Chakiachvili, M.; et al. Ensembl Genomes 2022: An Expanding Genome Resource for Non-Vertebrates. Nucleic Acids Research 2022, 50, D996–D1003, doi:10.1093/nar/gkab1007.

31. Higo, K.; Ugawa, Y.; Iwamoto, M.; Korenaga, T. Plant Cis-Acting Regulatory DNA Elements (PLACE) Database: 1999. Nucleic Acids Res. 1999, 27, 297–300.

32. Yilmaz, A.; Mejia-Guerra, M.K.; Kurz, K.; Liang, X.; Welch, L.; Grotewold, E. AGRIS: The Arabidopsis Gene Regulatory Information Server, an Update. Nucleic Acids Research 2011, 39, D1118–D1122, doi:10.1093/nar/gkq1120.

33. Castro-Mondragon, J.A.; Riudavets-Puig, R.; Rauluseviciute, I.; Berhanu Lemma, R.; Turchi, L.; Blanc-Mathieu, R.; Lucas, J.; Boddie, P.; Khan, A.; Manosalva Pérez, N.; et al. JASPAR 2022: The 9th Release of the Open-Access Database of Transcription Factor Binding Profiles. Nucleic Acids Res 2021, 50, D165–D173, doi:10.1093/nar/gkab1113.

34. Nguyen, N.T.T.; Contreras-Moreira, B.; Castro-Mondragon, J.A.; Santana-Garcia, W.; Ossio, R.; Robles-Espinoza, C.D.; Bahin, M.; Collombet, S.; Vincens, P.; Thieffry, D.; et al. RSAT 2018: Regulatory Sequence Analysis Tools 20th Anniversary. Nucleic Acids Research 2018, 46, W209–W214, doi:10.1093/nar/gky317.

